# Toxin-Triggered Liposomes for the Controlled Release of Antibiotics to Treat Infections Associated with Gram-Negative Bacteria

**DOI:** 10.1101/2023.09.28.559931

**Authors:** Ziang Li, Rani Baidoun, Angela C. Brown

## Abstract

Antibiotic resistance has become an urgent threat to health care in recent years. The use of drug delivery systems provides advantages over conventional administration of antibiotics and can slow the development of antibiotic resistance. In the current study, we developed a toxin-triggered liposomal antibiotic delivery system, in which the drug release is enabled by the leukotoxin (LtxA) produced by the Gram-negative pathogen, *Aggregatibacter actinomycetemcomitans*. LtxA has previously been shown to mediate membrane disruption by promoting a lipid phase change in nonlamellar lipids, such as 1,2-dioleoyl-*sn*-glycero-3- phosphoethanolamine-N-methyl (N-methyl-DOPE). In addition, LtxA has been observed to bind strongly and nearly irreversibly to membranes containing large amounts of cholesterol. Here, we designed a liposomal delivery system composed of N-methyl-DOPE and cholesterol to take advantage of these interactions. Specifically, we hypothesized that liposomes composed of N-methyl-DOPE and cholesterol, encapsulating antibiotics, would be sensitive to LtxA, enabling controlled antibiotic release. We observed that liposomes composed of N-methyl-DOPE were sensitive to the presence of low concentrations of LtxA, and cholesterol increased the extent and kinetics of content release. The liposomes were stable under various storage conditions for at least 7 days. Finally, we showed that antibiotic release occurs selectively in the presence of an LtxA-producing strain of *A. actinomycetemcomitans* but not in the presence of a non-LtxA- expressing strain. Together, these results demonstrate that the designed liposomal vehicle enables toxin-triggered delivery of antibiotics to LtxA-producing strains of *A. actinomycetemcomitans*.

---

Antibiotic resistance, primarily caused by the overuse and misuse of antibiotics, continues to pose a significant threat to the global public health, despite warnings about the problem from multiple agencies, including the United States Centers for Disease Control (CDC) and World Health Organization (WHO).^1,2^ Several novel alternatives to conventional antibiotics have emerged in recent years, such as endolysins, bacteriophage-derived enzymes, antimicrobial peptides, and bacteriocins.^3–6^ Although many of these molecules are promising, traditional antibiotics are still heavily relied upon in treating infections due to their wide availability, inexpensive cost, and proven safety. However, in recent years, not only has the discovery of new classes of antibiotics declined drastically, but the time interval between the introduction of new drugs and the emergence of resistance to them by pathogens has also significantly shortened.^7,8^ The very use of antibiotics causes a selective pressure that leads to resistance;^2,9,10^ as long as antibiotics are used in clinical settings, the emergence of drug-resistant bacteria is an inevitable problem. Gram-negative bacteria are particularly problematic due to their complex cell wall, which consists of two lipid membranes that effectively prevent internalization of most antibiotics.^11^ As a result, no new classes of antibiotics that are effective against Gram-negative bacteria have been discovered since the quinolone family was introduced in the 1970s.^8^ It is therefore imperative that we improve the administration of currently efficacious antibiotic agents to slow the development of antimicrobial resistance, while alternative treatments remain under development.^12^ The use of targeted delivery and controlled release systems is a promising strategy for antimicrobial treatments amid the antibiotic crisis, as the use of drug carriers for targeted delivery reduces the rate of acquired antibiotic resistance due to increased local concentration at the site of infection and decreased exposure in non-infected sites. Furthermore, reduced drug side effects and increased drug uptake can be achieved through the use of suitable antibiotic delivery systems.^12–16^

*Aggregatibacter actinomycetemcomitans* is an oral Gram-negative bacterium that has been closely associated with inflammatory periodontal disease, particularly aggressive forms of periodontitis.^17,18^ Without proper and timely treatments, these aggressive forms of periodontitis lead to the loss of connective and supporting tissue and damaged alveolar bone and eventually tooth loss, affecting patients physically and cosmetically.^18,19^ The traditional treatment for periodontitis is scaling and root planing (SRP), sometimes followed by periodontal flap surgery.^20^ However, these mechanical treatments are not always effective since they are unable to suppress bacterial infections at the inflammation sites, thus allowing *A. actinomycetemcomitans* to persist.^21–25^ As a result, antibacterial agents have been used systemically as a supplement to SRP to treat aggressive periodontitis. Although adjunctive antibiotic treatment has been reported to improve therapeutic outcomes,^26–28^ systemic antibiotics have drawbacks, including a risk for drug side effects^29^ and increased development of antibiotic resistance.^15,30^ In particular, subtherapeutic concentrations in the blood and at the infection site^31^ fail to eradicate the pathogen completely, contributing to the development of antibiotic-resistant species.^32^ Over the years, several patient studies have reported resistance of *A. actinomycetemcomitans* to numerous commonly used oral antimicrobial agents, including amoxicillin, tetracycline, doxycycline, and metronidazole. Therefore, there is an essential need for targeted delivery strategies to treat *A. actinomycetemcomitans*-associated periodontitis to decrease the risk of antibiotic resistance.^33–37^

*A. actinomycetemcomitans* possesses an arsenal of virulence factors that facilitate its pathogenesis and infection, one of which is a leukotoxin (LtxA), a membrane-disrupting toxin belonging to the repeats-in-toxin (RTX) family that specifically attacks human immune cells.^38–42^ Due to this immunosuppressive activity and the observation that highly leukotoxic strains of *A. actinomycetemcomitans*, such as JP2, are more closely associated with disease,^43^ the mechanism of action of LtxA has been studied in detail.^39,44–47^ After secretion by *A. actinomycetemcomitans* via a Type I secretion system,^44,45^ LtxA interacts specifically with immune cells via recognition of an integrin receptor, lymphocyte function-associated antigen-1 (LFA-1) on the cell surface.^39,46^ In addition, LtxA binds to cholesterol on the target cell membrane through a cholesterol recognition amino acid consensus (CRAC) domain.^47^

During its insertion into the plasma membrane, LtxA disrupts the integrity of the host cell membrane bilayer. A series of biophysical assays in liposomes demonstrated that LtxA-mediated content leakage occurred only when the liposome composition included lipids with a negative spontaneous curvature.^48^ These types of lipids are able to form nonlamellar phases,^49^ such as the inverted hexagonal (H_II_) phase. The incorporation of the bilayer-stabilizing lipid cholesterol sulfate inhibited LtxA-mediated membrane disruption.^48^ These results suggested that LtxA might mediate membrane disruption, not through pore formation, but via destabilization of the bilayer by promotion of nonlamellar phase formation. Subsequent work demonstrated that indeed, LtxA is able to promote the bilayer-to-H_II_ phase transition in liposomes containing lipids with negative spontaneous curvature, in particular, 1,2-dioleoyl-*sn*-glycero-3-phosphoethanolamine-N-methyl (N-methyl-DOPE).^48^

Because of the reported association of LtxA with disease severity, we have proposed the use of LtxA as a therapeutic target. Using our knowledge of the mechanisms by which LtxA recognizes and interacts with immune cell membranes, our lab has previously developed several LtxA-focused strategies that aimed to counteract the immunosuppressive and cytotoxic activity of LtxA. For example, we demonstrated that a peptide based on the CRAC sequence of LtxA was able to block the toxin’s recognition of cholesterol and inhibit LtxA-mediated activity.^50^ We have also designed peptides based on the domains of LFA-1 recognized by LtxA and showed their effectiveness in inhibiting LtxA cytotoxicity by disrupting LtxA-LFA-1 recognition.^51^ In addition to these two receptor-blocking approaches, we have also demonstrated that epigallocatechin gallate (EGCg), a type of catechin abundantly found in green tea extract, alters the secondary structure of LtxA, reducing its affinity for cholesterol on the host cell membrane^52^ and impacting the secretion and localization of LtxA.^53^

In this work, we were similarly motivated by the association of LtxA and the pathogenicity of *A. actinomycetemcomitans* and aimed to use LtxA as a trigger for controlled antibiotic release from a liposomal delivery vehicle. Based on the high affinity of LtxA for cholesterol and its ability to disrupt lipid membranes in a specific manner, we hypothesized that a liposome encapsulating antibiotics and composed of cholesterol and an H_II_-forming lipid would be disrupted only in the presence of LtxA (Fig. 1), that is, in the presence of disease-associated strains of *A. actinomycetemcomitans*. Our results demonstrate that this “Trojan Horse” delivery system is effective in releasing antibiotics in a controlled, toxin-dependent manner.

**Figure 1.**
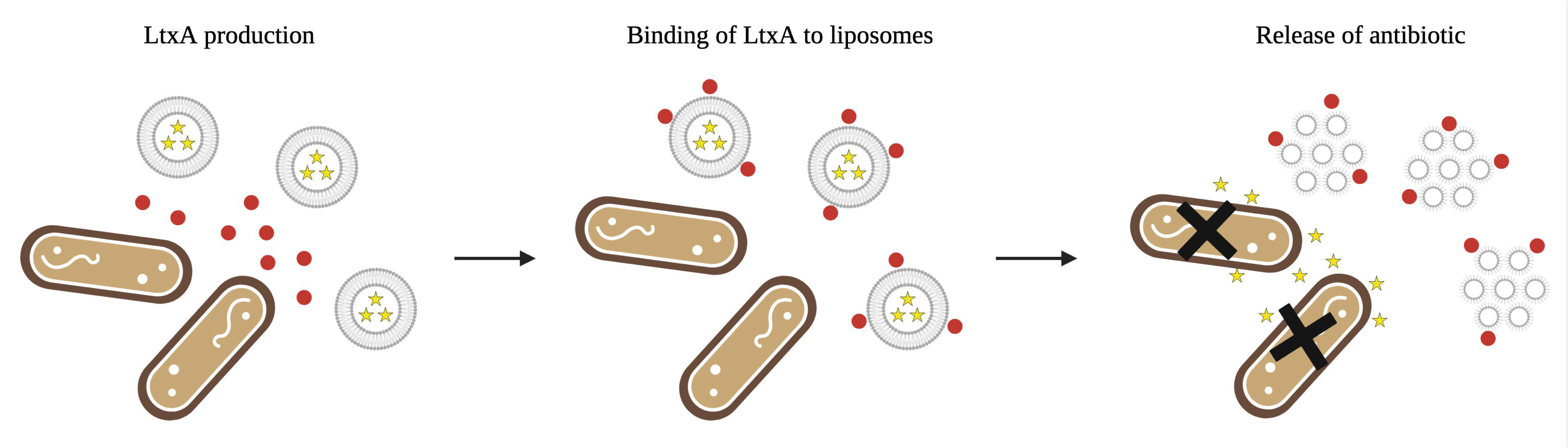
Mechanism of LtxA-triggered antibiotic release. LtxA is produced and secreted by *A. actinomycetemcomitans*. LtxA then binds to the antibiotic-loaded liposomes due to strong affinity for cholesterol. An LtxA-mediated phase transition occurs upon the interaction between LtxA and liposomes, causing leakage of encapsulated antibiotic which, consequently, leads to the death of *A. actinomycetemcomitans* cells. Figure created with BioRender.com.

## Results

### LtxA promotes content leakage from N-methyl-DOPE liposomes

Previous work has shown that LtxA promotes a bilayer-to-H_II_ transition in liposomes composed of N-methyl-DOPE.^48^ Here, we used a leakage assay to investigate whether this toxin-mediated phase transition is sufficient to cause content release from the liposomes. First, 8- aminonaphthalene-1,3,6-trisulfonic acid, disodium salt (ANTS)/ *p*-xylene-bis-pyridinium bromide (DPX)-loaded liposomes (250 µM) composed of 100% N-methyl-DOPE were incubated with LtxA (0.100, 0.125, and 0.167 µM) for 30 min, and the resulting leakage was quantified by measuring the emission intensity of ANTS and calculating a %Release using Eq. 1. As shown in Fig. 2A, liposomes composed of 100% N-methyl-DOPE were highly responsive to the presence of LtxA, with increasing toxin concentrations resulting in increased release of the fluorescent probe.

**Figure 2.**
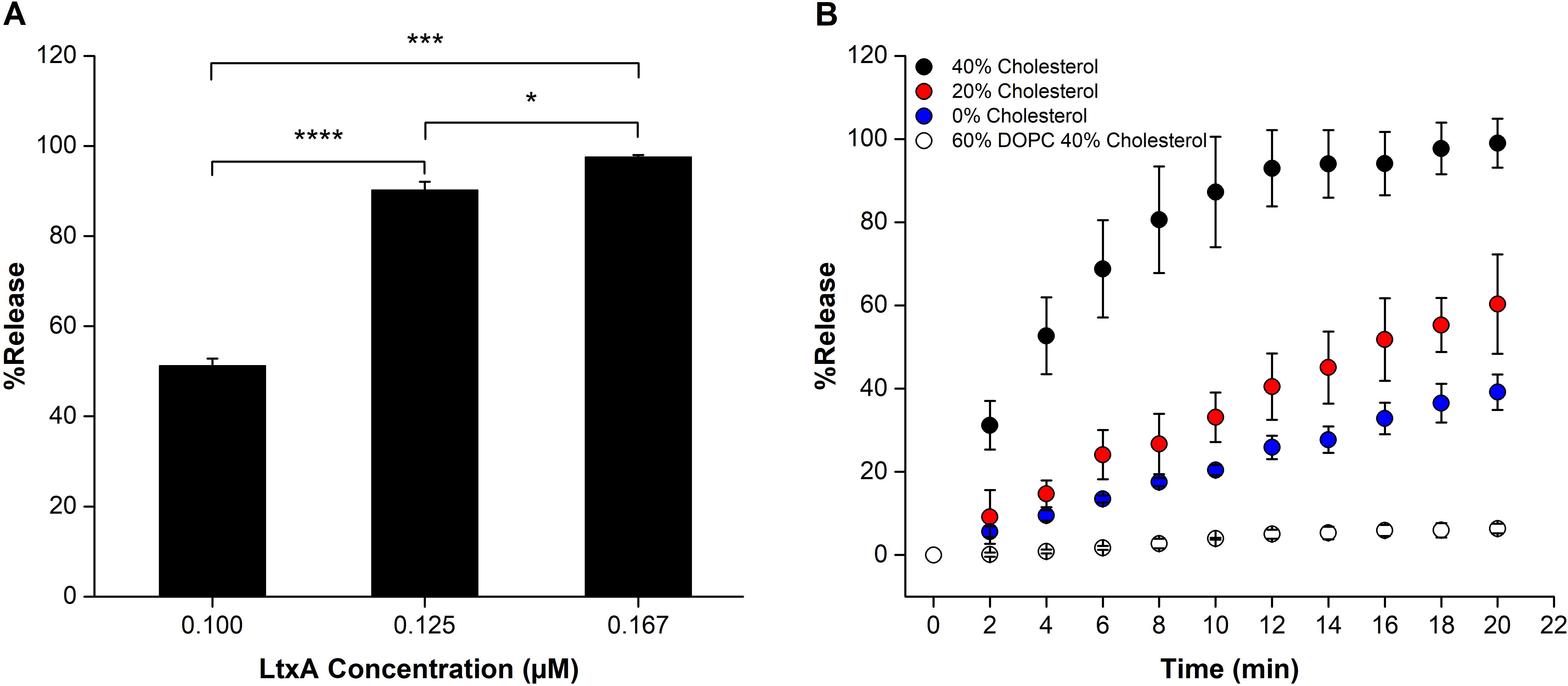
LtxA-mediated leakage from liposomes. (A) Liposome leakage as a function of LtxA concentration. N-methyl-DOPE liposomes (250 µM), encapsulating ANTS and DPX were treated with LtxA at different concentrations, and %Release was measured after 30 min. Each value represents the mean (n = 3) ± standard deviation. T-test results: *, p<0.05; ***, p<0.001; ****, p<0.0001. (B) ANTS/DPX release profiles from liposomes (250 µM) composed of N- methyl-DOPE with 0% (blue), 20% (red), or 40% (black) cholesterol and liposomes composed of DOPC and 40% cholesterol (white), each treated with 0.100 µM LtxA. The statistical significance was analyzed using ANOVA (Table S1).

We next investigated whether incorporation of cholesterol in the N-methyl-DOPE liposomes could enhance LtxA-mediated leakage from the liposomes. Using an LtxA concentration of 0.100 µM, we measured the rate of content release from liposomes composed of N-methyl-DOPE and 0%, 20%, or 40% cholesterol. Incorporation of 20% cholesterol in the liposomes increased the amount of leakage slightly relative to 100% N-methyl-DOPE liposomes, while incorporation of 40% cholesterol increased the rate and amount of leakage to an even greater extent (Fig. 2B). These trends demonstrate that LtxA-mediated leakage from N-methyl- DOPE liposomes can be improved by incorporating cholesterol into the liposomal membrane.

N-methyl-DOPE is a lipid that forms bilayers at room temperature but is able to transition to an H_II_ phase at temperatures above 55 °C.^54^ In contrast, 1,2-dioleoyl-*sn*-glycero-3- phosphocholine (DOPC) is unable to transition to an H_II_ phase at any temperature. Therefore, to demonstrate that the observed LtxA-mediated leakage is specifically the result of this bilayer-to- H_II_ phase transition, we repeated the experiment using liposomes composed of 60% DOPC and 40% cholesterol as a control. As presented in Fig. 2B, very little release was observed for DOPC/Cholesterol liposomes, demonstrating that the observed content release occurs only upon LtxA-mediated phase change. We chose to use liposomes composed of 60% N-methyl-DOPE/40% Chol for the remainder of the study because of the strong sensitivity of this composition to LtxA.

### N-methyl-DOPE liposomes are stable

To examine the stability of the N-methyl-DOPE liposomes, we measured their hydrodynamic diameter over time at 4°C and room temperature (∼22°C). Using dynamic light scattering (DLS), we observed that both empty and antibiotic-encapsulated liposomes remained stable under all three conditions throughout seven days in terms of their mean hydrodynamic particle diameter and polydispersity indices (PDIs, Fig. 3, Table S2).

**Figure 3.**
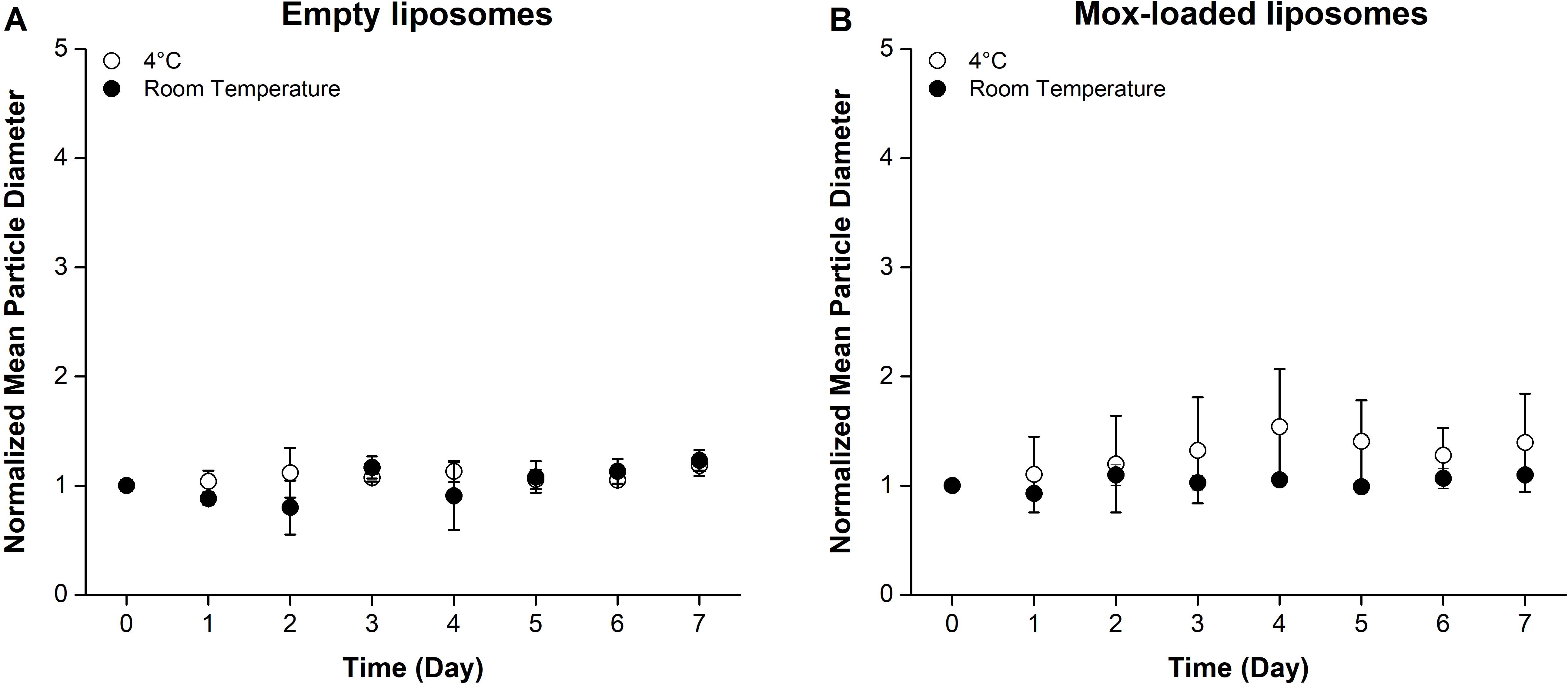
Mean size of N-methyl-DOPE liposomes over 7 consecutive days. The normalized mean hydrodynamic diameter of empty (A) and moxifloxacin-containing (B) liposomes composed of 60% N-methyl-DOPE and 40% cholesterol stored at 4°C and room temperature (∼22°C), as measured by DLS for 7 days. Each of the data points was normalized to the corresponding initial (Day 0) value. Each data point represents the mean (n=3) ± standard deviation.

### N-methyl-DOPE liposomes are able to encapsulate moxifloxacin

The minimum inhibitory concentration (MIC) of moxifloxacin for *A. actinomycetemcomitans* cells has been reported to be within the range of 0.006 to 0.032 µg/mL.^33,34,36,55^ To confirm the antimicrobial effect of moxifloxacin against the JP2 clone of *A. actinomycetemcomitans* in our lab setting, we treated *A. actinomycetemcomitans* JP2 cells with five different concentrations of moxifloxacin. The optical density measured at a wavelength of 600 nm (OD_600_) readings of the untreated and moxifloxacin-treated cells after a 24-hour incubation at 37°C were collected and plotted against the moxifloxacin concentration (Fig. S1A). The *A. actinomycetemcomitans* JP2 cells exhibited susceptibility to moxifloxacin at concentrations greater than 0.008 µg/mL, demonstrating strong antimicrobial effectiveness of moxifloxacin against these cells.

To quantify the encapsulation efficiency of moxifloxacin in the N-methyl-DOPE liposomes we used UV-vis spectrophotometry. As shown in Fig. S1B, moxifloxacin exhibited significant absorption at a wavelength of approximately 335 nm, and the peak increased proportionally with the moxifloxacin concentration. A calibration curve consisting of the absorbance at 335 nm for varying moxifloxacin concentrations was then constructed (Fig. 4). Following the separation of unencapsulated moxifloxacin from antibiotic-loaded liposomes using centrifugal filtration, the unencapsulated drug was serially diluted, and the absorbance at 335 nm was measured. Using the calibration curve, the encapsulation efficiency was calculated to be 57.8% ± 1.2%.

**Figure 4.**
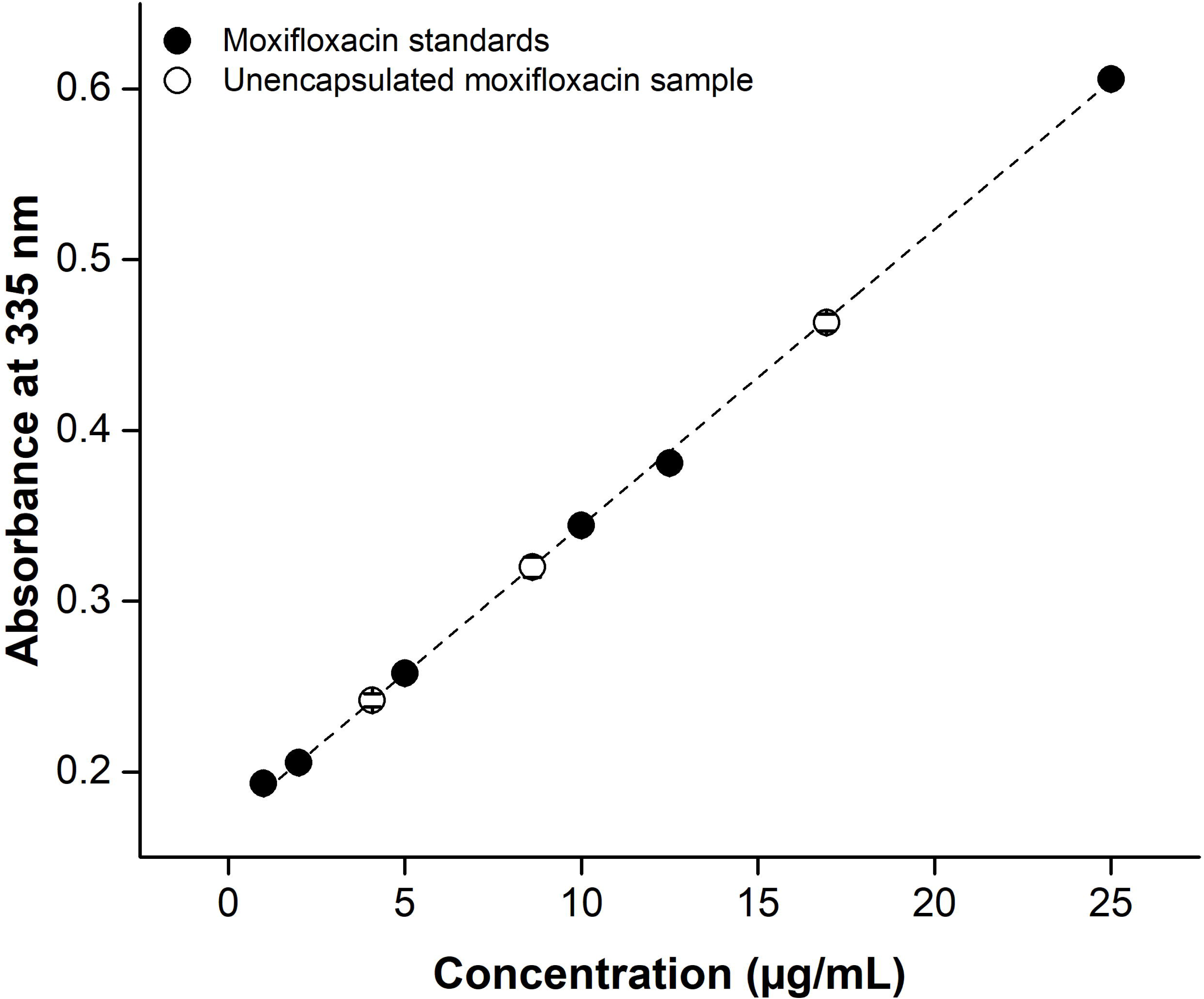
Calculation of Encapsulation Efficiency. The absorbance at 335 nm of moxifloxacin solutions at known concentrations (black) was measured and used to create a calibration curve. Unencapsulated moxifloxacin was removed from the liposomes using centrifugal filtration, and the absorbance of the serially diluted filtrate (white) was plotted on the calibration curve to calculate the mass of unencapsulated moxifloxacin. Each data point represents the mean (n=3) ± standard deviation.

### N-methyl-DOPE liposomes selectively release moxifloxacin in the presence of LtxA-expressing strains of *A. actinomycetemcomitans*

To evaluate the effectiveness of the moxifloxacin-encapsulating N-methyl-DOPE/Chol liposomes against *A. actinomycetemcomitans* cells, we compared the activity of the liposomes against two strains of *A. actinomycetemcomitans*: JP2, an LtxA-producing strain and AA1704, an isogenic mutant that does not express LtxA.^56^ Both cell types were treated with either (1) free moxifloxacin or (2) moxifloxacin-loaded liposomes, and the OD_600_ readings were recorded as a measure of cell viability. The increase in OD_600_ due to the presence of liposomes in treatment (2) was accounted for by subtracting the signal difference between cells with no treatment and cells treated with unloaded liposomes at the same lipid concentration as treatment (2).

As illustrated in Fig. 5A, both strains were first grown for 6 hr to a time point where JP2 cells begin to secrete LtxA, which was confirmed by an immunoblot test (Fig. S2). After this initial 6-hr incubation, the cells were grown for an additional 4 hr, either untreated or treated with one of the two treatments (free moxifloxacin or liposome-encapsulated moxifloxacin, both with the same moxifloxacin concentration). As expected, the growth of both JP2 (Fig. 5B) and AA1704 cells (Fig. 5C) was greatly inhibited by free moxifloxacin, at each time point. The growth of the JP2 cells was also significantly inhibited in the presence of the moxifloxacin- loaded liposomes. In contrast, the growth of AA1704 cells, which do not express LtxA, was only slightly inhibited by the moxifloxacin-loaded liposomes.

**Figure 5.**
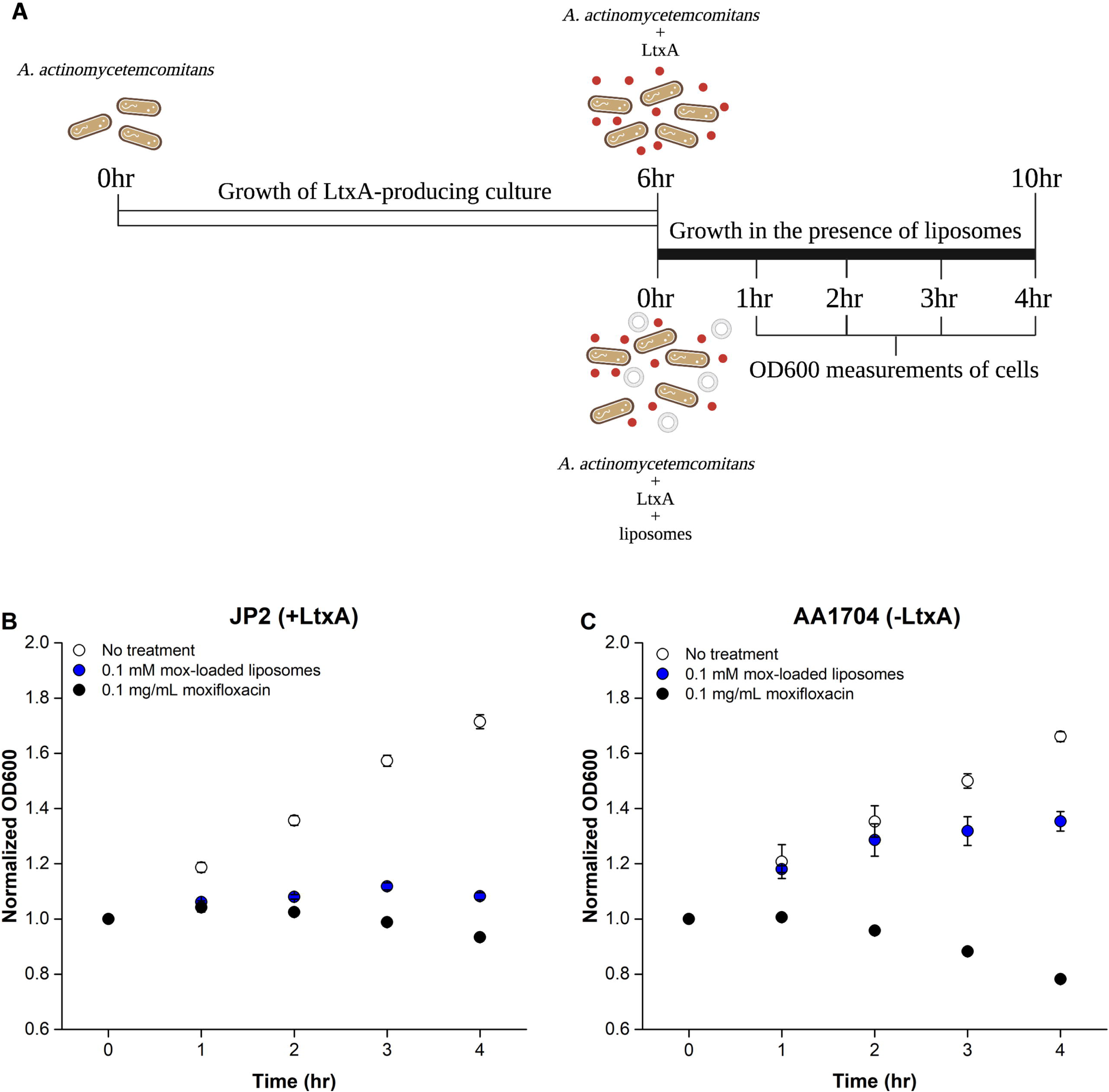
**Growth of *A. actinomycetemcomitans* cells in the presence of different treatments.** (A) Timeline of *A. actinomycetemcomitans* cell growth. *A. actinomycetemcomitans* cells were grown for 6 hr to the point where LtxA secretion is initiated, followed by a 4-hr growth in the presence of the liposome treatments. Created with BioRender.com. The plots show the growth of (B) *A. actinomycetemcomitans* JP2 cells (which express LtxA) or (C) *A. actinomycetemcomitans* AA1074 cells (which do not express LtxA) in the absence of treatment (white), or after treatment with moxifloxacin solution at 0.1 mg/mL (black) or moxifloxacin-loaded N-methyl-DOPE liposomes at 0.1 mM (blue), normalized by the initial OD_600_ value. Each data point represents the average (n=3) ± standard deviation. The statistical significance was analyzed using ANOVA (Table S3).

To better compare the specific inhibitory action of the moxifloxacin-loaded N-methyl- DOPE/Chol liposomes against the two strains of *A. actinomycetemcomitans*, the percent inhibition was calculated (Eq. 2) at each time point for both *A. actinomycetemcomitans* JP2 and AA1704. As shown in Fig. 6, antibiotic-loaded liposomes exhibited great bactericidal effectiveness against the LtxA-expressing JP2 cells at all time points. The moxifloxacin-loaded liposomes inhibited the growth of the LtxA-expressing JP2 cells more than twice as effectively as the non-LtxA-expressing AA1704 cells at all timepoints. Together, these results demonstrate the specificity of LtxA in mediating the release of moxifloxacin from the N-methyl-DOPE liposomes and indicate the suitability of this formulation for the targeted release of antibiotics due to LtxA.

**Figure 6.**
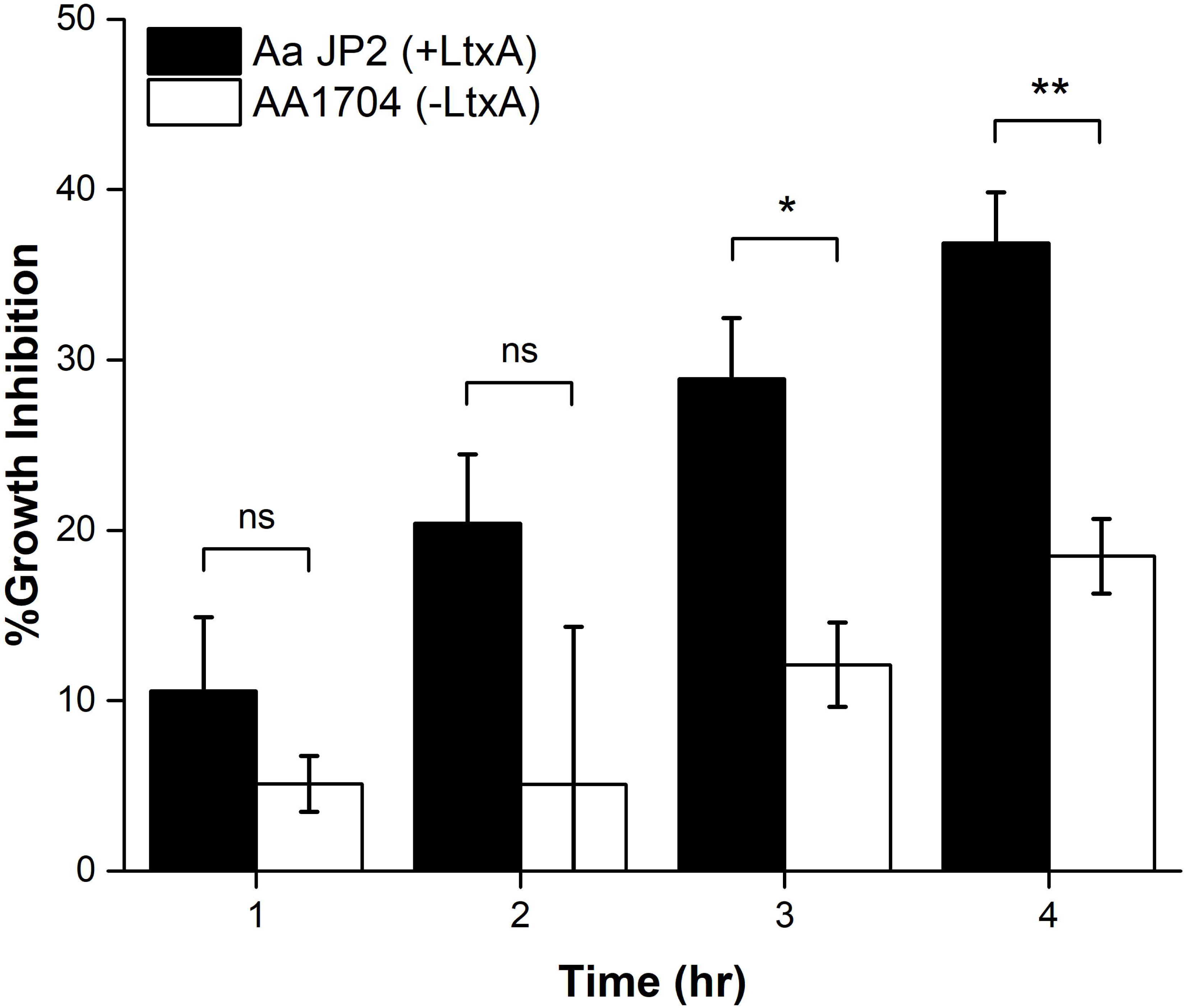
Inhibition of *A. actinomycetemcomitans* growth by moxifloxacin-loaded liposomes. Inhibition of growth of JP2 (black) and AA1704 (white) cells after treatment with moxifloxacin- loaded liposomes. Each data point represents the mean (n=3) ± standard deviation. T-test results: ns, not significant; *, p<0.05; **, p<0.01.

## Discussion

SRP is the traditional method of treating *A. actinomycetemcomitans* infections, but in many cases, this approach is insufficient in completely clearing the infection.^22,25^ Therefore, SRP is commonly applied in combination with systemically administered antibiotics to improve therapeutic outcomes.^57,58^ Nonetheless, prolonged use of systemic antibiotics increases the risk of development of antibiotic resistance in periodontal pathogens; short-term and intermittent use of antibiotics can also lead to drug resistance, making antimicrobial treatment less effective.^59^ For example, in recent years, many *A. actinomycetemcomitans* isolates have been observed to be less susceptible to amoxicillin, azithromycin, metronidazole and tetracycline, the antibiotic agents frequently used in periodontal treatments.^36^

The use of delivery systems for the controlled release of antibiotics can slow the development of drug resistance by increasing the local drug concentration and minimizing exposure of the microbiota to the drugs.^12,14,60^ In this work, we therefore sought to develop a delivery vehicle that releases antibiotics only in the presence of pathogenic strains of *A. actinomycetemcomitans*. To accomplish this, we focused on LtxA, as its expression has been correlated with disease progression.^61–63^ We utilized the unique mechanism of LtxA-mediated membrane disruption, which occurs through the induction of a bilayer-to-H_II_ phase transition^48^ to design a liposomal delivery vehicle that employs LtxA as a trigger for the controlled delivery of antibiotics to *A. actinomycetemcomitans*. Our results demonstrated that the LtxA-mediated phase transition was adequate to enable antibiotic release from N-methyl-DOPE liposomes and that the integration of cholesterol greatly promoted toxin-triggered release. More importantly, we demonstrated that most of the antibiotic release was specifically caused by the presence of LtxA. These results support our hypothesis that this liposome formulation is a promising toxin-triggered and targeted antibiotic delivery vehicle for the treatment of *A. actinomycetemcomitans* infections.

Liposomes have been extensively explored as a drug delivery carrier thanks to their biocompatibility, excellent tunability in terms of size and composition, and ability to carry both hydrophilic and hydrophobic molecules.^14^ Prolonged circulation time can be achieved by incorporating polyethylene glycol in the liposomal compositions,^14^ which improves antibiotic efficacy while reducing dosing frequency and the required concentration of drugs.^64^ With the wide variety of available lipids from which to construct liposomes, numerous responsive liposomal drug delivery vehicles have been developed to respond to external stimuli, such as pH, light, ultrasound, the presence of specific proteins including enzymes and toxins.^65,66^

A distinguishing feature of our delivery vehicle is its sensitivity to a Gram-negative bacterial toxin. Prior work to develop toxin-responsive liposomes have focused on the staphylococcal α-toxin, which produces large, protein lined pores in a non-specific manner.^67,68^ Pornpattananangkul, et al. reported a gold-chitosan-stabilized, cholesterol-containing liposomal system that released encapsulated vancomycin upon pore formation by the staphylococcal α- toxin.^69^ More recently, Yang et al. coated mesoporous silica nanoparticles with a lipid bilayer and investigated their potential for intracellular antibiotic release by α-toxin; the authors showed that the intracellular delivery of gentamicin was improved by conjugating the lipid nanoparticles with a targeting ligand derived from an antimicrobial peptide.^70^ Finally, Wu et al., developed a toxin-responsive “nanoreactor” by encapsulating calcium peroxide and rifampicin in a fatty acid core with a lipid monolayer. Pore formation by the staphylococcal α-toxin enabled the penetration of water into the core, where it reacted with the calcium peroxide to promote antibiotic release.

In contrast to these published works, we have, here, focused on the unique interaction of LtxA with non-lamellar lipids that enables a phase change (rather than classical pore formation), and demonstrated that this approach is sufficient to promote toxin-responsive antibiotic release resulting in bacterial death. LtxA, and the RTX family of proteins, to which it belongs,^39,45,46,71^ have long been described as “pore-forming toxins” due to their ability to induce a large increase in the cytosolic calcium concentration of target cells.^72^ However, prior work demonstrated that LtxA does not form a pore in the classical sense, but rather destabilizes bilayer packing through the mediation of a lipid phase change.^48^ Among the multiple pieces of evidence supporting the LtxA-mediated phase change was a series of liposome leakage experiments that demonstrated that LtxA was only able to disrupt the lipid bilayer when the membrane contained nonlamellar lipids, such as phosphatidylethanolamine (PE).^48^ Despite differences in receptor specificity, many RTX toxins share this ability to disrupt cell membranes containing nonlamellar lipids,^73–75^ suggesting that the mechanism of toxin-mediated phase change is shared among these toxins. For example, the *Escherichia coli* α-hemolysin (HlyA) is able to induce membrane leakage from liposomes composed of phosphatidylcholine (PC), PE, and cholesterol, with the incorporation of a PE lipid in the liposomes enhancing HlyA-triggered release.^76^ This common behavior suggests that our proposed antibiotic delivery system may also be applicable in the targeted treatment of other pathogenic Gram-negative bacteria that produce RTX toxins.

In addition to this shared process of membrane disruption, LtxA and many of the RTX toxins possess a CRAC domain that enables strong binding to cholesterol.^47,77–81^ For this reason, we incorporated a large proportion of cholesterol in our liposomal formulation and observed that the incorporation of cholesterol in the liposomes greatly enhanced LtxA-mediated leakage. Prior work has used a similar approach, using natural or synthetic cell membranes to sequester toxins produced by Gram-positive bacteria due to their affinity for specific lipid components. For example, a toxin-binding “nanosponge” system consisting of a poly(lactic-*co*-glycolic acid) core coated with a red blood cell membrane was shown to neutralize several toxins, including the staphylococcal α-toxin, streptolysin O and melittin due to the strong affinity of these toxins for the red blood cell membrane.^82^ In a similar approach, Jiang et al. created a hybrid liposome composed of natural erythrocyte membranes and polymyxin B-modified lipids. The polymyxin B provided affinity for the bacterial cell membrane, while the red blood cell membrane acted to absorb toxins released by the bacteria.^83^ A fully synthetic toxin-absorbing liposome, composed of sphingomyelin and cholesterol, was developed by Henry et al. and effectively sequestered multiple toxins secreted by Gram-positive bacteria. The *in vivo* studies demonstrated that the bacteria were eliminated by the host’s immune system following successful protection from bacterial toxins by the liposomes.^84^

Importantly, none of these liposomal delivery systems is responsive to the presence of toxins released by Gram-negative bacteria. This class of bacteria is particularly refractory to antibiotics, due to their intrinsic resistance caused by their dual-membrane cell wall as well as acquired mechanisms of resistance including expression of antibiotic-degrading enzymes and/or efflux pumps.^85^ With the lack of suitable antibiotics to treat infections associated with Gram-negative bacteria, new treatment options, such as the one described here, are urgently needed. Our liposomal antibiotic delivery vehicle combines the sequestration ability of cholesterol-containing membranes and uses the unique mechanism of toxin-mediated membrane disruption employed by the RTX toxins to enable controlled antibiotic release. Because the RTX family of toxins represents one of the largest group of toxins produced by Gram-negative bacteria,^85^ we anticipate that the approach developed here will have broad applicability for the treatment of Gram-negative infections.

## Conclusion

The current work demonstrates the potential of a novel toxin-triggered liposomal antibiotic delivery vehicle for the targeted treatment of *A. actinomycetemcomitans* infections. This formulation takes advantage of the strong affinity of LtxA for cholesterol and the ability of LtxA to disrupt the membrane by mediating a membrane phase change. Due to significant similarities among RTX toxins in their ability to disrupt lipid membranes in this unique manner, we expect that a similar approach could be broadly applied to infections caused by other RTX toxin-producing Gram-negative pathogens.

## Materials and Methods

### Chemicals

ANTS and DPX were purchased from ThermoFisher Scientific. N-methyl-DOPE was purchased from Avanti Lipids (Alabaster, AL). Cholesterol was purchased from Sigma-Aldrich. All reagents were used without additional purification.

### Bacterial Strains and Cell Cultures

Two strains of *A. actinomycetemcomitans*, JP2 and AA1704, were used in the current study.^57^ Both JP2 and AA1704 were inoculated and anaerobically cultured in *A. actinomycetemcomitans* growth medium (AAGM)^86^ in a candle jar at 37 °C for 24 hr. The bacteria were then transferred to a larger AAGM culture, in which they were grown in an incubator at 37 °C.

### LtxA Purification

*A. actinomycetemcomitans* JP2 cells were pelleted in a centrifuge at 10,000 xg for 10 min at 4 °C following 24-hr incubation of the cells in 1 L of AAGM at 37°C. The cell pellets were then discarded, and LtxA was purified from the culture supernatant following a published procedure.^87^ The purity of the purified LtxA was determined by SDS-PAGE and its identity was confirmed using a western blot.

### Preparation of Liposomes

N-methyl-DOPE (25 mg/mL in chloroform) and cholesterol (25 mg/mL in chloroform) were added to a glass vial at the desired ratios. The chloroform was then evaporated inside a fume hood using nitrogen flow to leave a thin layer of lipid film on the vial interior. The lipid film was placed under vacuum overnight to remove residual chloroform. Two separate samples were simultaneously prepared in the same manner. One was hydrated with HEPES buffer (50 mM NaCl, 25 mM HEPES, pH 7.4) and later used as a standard solution to determine the concentration of loaded liposomes. The other was hydrated with an ANTS/DPX buffer (12.5 mM ANTS, 45 mM DPX, 50 mM NaCl, 25 mM HEPES, pH 7.4) or a moxifloxacin solution (10 mg/mL) depending on the type of tests in which the liposomes were used. The lipid films were uniformly suspended by vortexing and were then mechanically extruded 15 times through 100 nm polycarbonate filters to make unilamellar liposomes. Unencapsulated cargo was removed either using size exclusion chromatography (SEC) or by centrifugal filtration. For SEC, the liposome suspension was run through a chromatography column packed with Sephadex G-50 gel. Liposomes were eluted using HEPES buffer and were collected in Eppendorf microcentrifuge tubes. The empty liposomes were serially diluted and used as standards to construct a calibration curve of absorbances at 600 nm vs liposome concentrations in UV-vis spectrophotometer. Each liposome-containing fraction was combined and was then measured in UV-vis spectrophotometer to determine the final concentration after SEC. For centrifugal filtration, the liposome suspension was placed into an Amicon® Ultra-0.5, 3,000Da molecular weight cutoff centrifugal filter and centrifuged at 10,000 xg for 15 min.

### LtxA-induced leakage from N-methyl-DOPE liposomes

Liposomes loaded with ANTS/DPX were prepared using 100% N-methyl-DOPE according to the procedure described above and were diluted to a concentration of 0.25 mM with HEPES buffer. While the concentration of liposomes was fixed, LtxA in the same volume at 0.167, 0.125, and 0.100 µM was added to obtain lipid-to-toxin molar ratios of 1500, 2000, and 2500, respectively. Following a 30 min incubation at room temperature, a 50 µL aliquot of sample at each ratio was added into a quartz cuvette. The release of ANTS from liposomes was measured using Photon Technology International Quantamaster 400 fluorescence spectrometer (excitation at 350 nm, emission scan from 450 nm to 600 nm, emission peak at 500 nm), and the fluorescence intensity at 500 nm was recorded as I_S_. The baseline signal (I_B_) was measured using liposomes alone. The complete release of encapsulated ANTS was then obtained by adding 10 µL of 2.5% Triton X-100 to the same liposome sample, and the intensity at 500 nm was recorded as I_T_. The percentage of LtxA-induced release at different toxin concentrations was then calculated using the following equation:

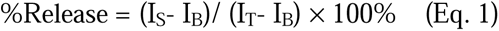

### Moxifloxacin encapsulation efficiency

A stock solution of moxifloxacin at 10 mg/mL was prepared by dissolving 150 mg moxifloxacin HCl solid in 10 mL of HEPES buffer solution. This stock solution was diluted to a secondary stock solution at a concentration of 50 µg/mL. The secondary stock was used to obtain the UV-vis absorption spectrum of moxifloxacin from 250-400 nm, and then was serially diluted to 6 different concentrations: 25, 12.5, 10, 5, 2 and 1 µg/mL. The absorbance of each diluted solution was measured in a Tecan plate reader and plotted against the concentrations to construct a calibration curve.

To determine the encapsulation efficiency of moxifloxacin in the N-methyl-DOPE liposomes, the concentration of unencapsulated moxifloxacin was determined first and used to calculate the amount of encapsulated drug. After separation of the moxifloxacin-loaded liposomes and unencapsulated moxifloxacin, as described above, the filtrates containing unloaded drug were then collected, consolidated and serially diluted before their absorbance was measured in the plate reader. The concentrations of the dilutions whose absorbance fell within the calibration curve range were calculated by fitting the absorbance values to the calibration equation. The encapsulation efficiency was then calculated by dividing the difference of the total mass of moxifloxacin and the mass of unencapsulated moxifloxacin, which yielded the amount of liposome-encapsulated drug, by the total mass of moxifloxacin.

### LtxA Immunoblotting

Immunoblotting was used to determine the time point at which LtxA production is initiated in *A. actinomycetemcomitans* JP2 cell culture. The starter culture was diluted with AAGM to an OD_600_ of 0.1, and a 1 mL aliquot was extracted and centrifuged at 10,000 xg for 10 min. The supernatant was reserved and stored at 4°C. This process was repeated every 1 hr for 8 hr. 2 µL of purified LtxA in Tris-buffered saline (TBS) (0.198M Tris HCl and 1.5M NaCl, pH 8.0) at six known concentrations (0.03125 µM, 0.0625 µM, 0.125 µM, 0.250 µM, 0.500 µM, and 1 µM) were blotted onto a nitrocellulose membrane as standards, and blotting of the reserved supernatant from each hour was performed on the same nitrocellulose membrane. After the membrane was dried, it was incubated in Blotto solution (5% powdered milk in TBS with 0.1% Tween 20 (TBST)) for 1 hr to block nonspecific antibody binding. The membrane was then incubated in a primary monoclonal antibody^88^ overnight, followed by 1 hr incubation with horseradish peroxidase-labeled goat anti-mouse secondary antibody (GAM-HRP, SouthernBiotech Cat#1010-05, RRID: AB_2728714). After washing, the membrane was incubated with SuperSignal™ West Dura Extended Duration Substrate (ThermoFisher™ Scientific) before being imaged using a BioRad ChemiDoc imaging system. The dot intensities were quantified using ImageJ.^89,90^ The concentrations of LtxA produced at each hour were determined using a calibration curve created with standard LtxA concentrations and corresponding intensities.

### Cell growth inhibition by encapsulated moxifloxacin

Moxifloxacin-loaded liposomes were prepared by following the procedure described above. Diluted starter cultures (OD_600_=0.1) of both JP2 and AA1704 strains were allowed to grow for 6 hr until the point of LtxA production. 100 µL of each of three different treatments were mixed into 900 µL of bacterial cells: (1) empty liposomes (final concentration in the samples, 0.1 mM), (2) moxifloxacin-encapsulating liposomes (final concentration, 0.1 mM), (3) moxifloxacin solution (final concentration, 0.1 mg/mL). A sample containing cells and HEPES buffer solution was prepared to monitor full growth of the bacterial cells, serving as an untreated control. The OD_600_ values of all samples were measured using a UV-vis spectrophotometer immediately after preparation and were recorded as the 0-hr readings. The samples were then incubated at 37°C and were read every hour in the spectrophotometer for 4 hr. All OD_600_ values were normalized by the corresponding 0-hr readings, and bacterial growth inhibition at each time point in the presence of different treatments was quantified by the equation below:

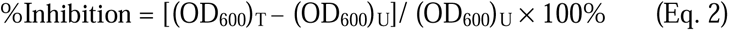

where (OD_600_)_T_ represents the OD_600_ values of the treated sample, and (OD_600_)_U_ represents the untreated sample.

## Supporting Information

Supporting information includes statistical analyses, hydrodynamic radii and PDI data, MIC determination, moxifloxacin UV-vis spectra, and LtxA immunoblot.

## Acknowledgments

The authors acknowledge funding from the National Institutes of Health (R21 DE032153) and Lehigh University (Faculty Innovation Grant program).

## Abbreviations Used

Mox: moxifloxacin
N-methyl-DOPE: 1,2-dioleoyl-*sn*-glycero-3-phosphoethanolamine-N- methyl
H_II_ phase: inverted hexagonal phase
DOPC: 1,2-dioleoyl-*sn*-glycero-3-phosphocholine
TBS: Tris-buffered saline
GAM-HRP: horseradish peroxidase-labeled goat anti-mouse
SEC: size-exclusion chromatography
AAGM: *Aggregatibacter actinomycetemcomitans* growth medium
SDS-PAGE: sodium dodecyl sulfate-polyacrylamide gel electrophoresis
SRP: scaling and planing
RTX: repeats-in-toxin
CRAC: cholesterol recognition amino acid consensus
EGCg: epigallocatechin gallate
ANTS: 8-aminonaphthalene-1,3,6-trisulfonic acid, disodium salt
DPX: *p*-xylene-bis-pyridinium bromide
DLS: dynamic light scattering
PDI: polydispersity index
MIC: minimum inhibitory concentration
OD_600_: optical density at 600 nm
HlyA: α-hemolysin

## References

1. World Health Organization. WHO Library Cataloguing-in-Publication Data Global Action Plan on Antimicrobial Resistance; 2015.

2. US Centers for Disease Control. Antibiotic Resistance Threats in the United States, 2019. 10.15620/cdc:82532.

3. Browne, K.; Chakraborty, S.; Chen, R.; Willcox, M. D. P.; Black, D. S.; Walsh, W. R.; Kumar, N. A New Era of Antibiotics: The Clinical Potential of Antimicrobial Peptides. International Journal of Molecular Sciences 2020, Vol. 21, Page 7047 2020, 21 (19), 7047. 10.3390/IJMS21197047.

4. Mahlapuu, M.; Håkansson, J.; Ringstad, L.; Björn, C. Antimicrobial Peptides: An Emerging Category of Therapeutic Agents. Front Cell Infect Microbiol 2016, 6 (DEC), 235805. 10.3389/FCIMB.2016.00194/BIBTEX.

5. Ghosh, C.; Sarkar, P.; Issa, R.; Haldar, J. Alternatives to Conventional Antibiotics in the Era of Antimicrobial Resistance. Trends Microbiol 2019, 27 (4), 323–338. 10.1016/J.TIM.2018.12.010.

6. Gondil, V. S.; Harjai, K.; Chhibber, S. Endolysins as Emerging Alternative Therapeutic Agents to Counter Drug-Resistant Infections. Int J Antimicrob Agents 2020, 55 (2), 105844. 10.1016/J.IJANTIMICAG.2019.11.001.

7. Silver, L. L. Challenges of Antibacterial Discovery. Clin Microbiol Rev 2011, 24 (1), 71–109. 10.1128/CMR.00030-10.

8. Lewis, K. The Science of Antibiotic Discovery. Cell 2020, 181 (1), 29–45. 10.1016/J.CELL.2020.02.056.

9. Hoffman, S. B. Mechanisms of Antibiotic Resistance. Compendium on Continuing Education for the Practicing Veterinarian 2001, 23 (5), 464–472. 10.1128/microbiolspec.vmbf-0016-2015.

10. Davies, J. Origins and Evolution of Antibiotic Resistance. Microbiologia 1996, 12 (1), 9–16. 10.1128/MMBR.00016-10.

11. Hancock, R. E. W. The Bacterial Outer Membrane as a Drug Barrier. Trends Microbiol 1997, 5 (1), 37–42. 10.1016/S0966-842X(97)81773-8.

12. Lai, C. K. C.; Ng, R. W. Y.; Leung, S. S. Y.; Hui, M.; Ip, M. Overcoming the Rising Incidence and Evolving Mechanisms of Antibiotic Resistance by Novel Drug Delivery Approaches – An Overview. Adv Drug Deliv Rev 2022, 181, 114078. 10.1016/J.ADDR.2021.114078.

13. Zhu, X.; Radovic-Moreno, A. F.; Wu, J.; Langer, R.; Shi, J. Nanomedicine in the Management of Microbial Infection – Overview and Perspectives. Nano Today 2014, 9 (4), 478–498. 10.1016/J.NANTOD.2014.06.003.

14. Huh, A. J.; Kwon, Y. J. “Nanoantibiotics”: A New Paradigm for Treating Infectious Diseases Using Nanomaterials in the Antibiotics Resistant Era. Journal of Controlled Release 2011, 156 (2), 128–145. 10.1016/J.JCONREL.2011.07.002.

15. Kalhapure, R. S.; Suleman, N.; Mocktar, C.; Seedat, N.; Govender, T. Nanoengineered Drug Delivery Systems for Enhancing Antibiotic Therapy. J Pharm Sci 2015, 104 (3), 872–905. 10.1002/JPS.24298.

16. Zhang, L.; Pornpattananangkul, D.; Hu, C.-M.; Huang, C.-M. Development of Nanoparticles for Antimicrobial Drug Delivery. Curr Med Chem 2010, 17 (6), 585–594. 10.2174/092986710790416290.

17. Fine, D. H.; Markowitz, K.; Furgang, D.; Fairlie, K.; Ferrandiz, J.; Nasri, C.; McKiernan, M.; Gunsolley, J. *Aggregatibacter Actinomycetemcomitans* and Its Relationship to Initiation of Localized Aggressive Periodontitis: Longitudinal Cohort Study of Initially Healthy Adolescents. J Clin Microbiol 2007, 45 (12), 3859–3869. 10.1128/JCM.00653-07.

18. Pihlstrom, B. L.; Michalowicz, B. S.; Johnson, N. W. Periodontal Diseases. The Lancet 2005, 366 (9499), 1809–1820. 10.1016/S0140-6736(05)67728-8.

19. Åberg, C. H.; Kelk, P.; Johansson, A. *Aggregatibacter Actinomycetemcomitans*: Virulence of Its Leukotoxin and Association with Aggressive Periodontitis. Virulence 2015, 6 (3), 188. 10.4161/21505594.2014.982428.

20. Sheridan, R. A.; Wang, H.-L.; Eber, R.; Oh, T.-J. Systemic Chemotherapeutic Agents as Adjunctive Periodontal Therapy- A Narrative Review and Suggested Clinical Recommendations. J Int Acad Periodontol 2015, 17 (4), 123–134.

21. Renvert, S.; Wikström, M.; Dahlén, G.; Slots, J.; Egelberg, J. Effect of Root Debridement on the Elimination of *Actinobacillus Actinomycetemcomitans* and *Bacteroides Gingivalis* from Periodontal Pockets. J Clin Periodontol 1990, 17 (6), 345–350. 10.1111/J.1600-051X.1990.TB00029.X.

22. Slots, J.; Rosling, B. G. Suppression of the Periodontopathic Microflora in Localized Juvenile Periodontitis by Systemic Tetracycline. J Clin Periodontol 1983, 10 (5), 465–486. 10.1111/J.1600-051X.1983.TB02179.X.

23. Mombelli, A.; Schmid, B.; Rutar, A.; Lang, N. P. Persistence Patterns of *Porphyromonas Gingivalis, Prevotella Intermedia*/ *Nigrescens*, and *Actinobacillus Actinomycetemcomitans* After Mechanical Therapy of Periodontal Disease. J Periodontol 2000, 71 (1), 14–21. 10.1902/JOP.2000.71.1.14.

24. Christersson, L. A.; Slots, J.; Rosling, B. G.; Genco, R. J. Microbiological and Clinical Effects of Surgical Treatment of Localized Juvenile Periodontitis. J Clin Periodontol 1985, 12 (6), 465–476. 10.1111/J.1600-051X.1985.TB01382.X.

25. Kornman, K. S.; Robertson, P. B. Clinical and Microbiological Evaluation of Therapy for Juvenile Periodontitis. J Periodontol 1985, 56 (8), 443–446. 10.1902/JOP.1985.56.8.443.

26. Soares, G. M. S.; Mendes, J. A. V.; Silva, M. P.; Faveri, M.; Teles, R.; Socransky, S. S.; Wang, X.; Figueiredo, L. C.; Feres, M. Metronidazole Alone or with Amoxicillin as Adjuncts to Non-Surgical Treatment of Chronic Periodontitis: A Secondary Analysis of Microbiological Results from a Randomized Clinical Trial. J Clin Periodontol 2014, 41 (4), 366–376. 10.1111/JCPE.12217.

27. Feres, M.; Haffajee, A. D.; Allard, K.; Som, S.; Socransky, S. S. Change in Subgingival Microbial Profiles in Adult Periodontitis Subjects Receiving Either Systemically-Administered Amoxicillin or Metronidazole. J Clin Periodontol 2001, 28 (7), 597–609. 10.1034/J.1600-051X.2001.028007597.X.

28. Rabelo, C. C.; Feres, M.; Gonçalves, C.; Figueiredo, L. C.; Faveri, M.; Tu, Y. K.; Chambrone, L. Systemic Antibiotics in the Treatment of Aggressive Periodontitis. A Systematic Review and a Bayesian Network Meta-Analysis. J Clin Periodontol 2015, 42 (7), 647–657. 10.1111/JCPE.12427.

29. Walker, C. B. Selected Antimicrobial Agents: Mechanisms of Action, Side Effects and Drug Interactions. Periodontol 2000 1996, 10 (1), 12–28. 10.1111/J.1600-0757.1996.TB00066.X.

30. Drulis-Kawa, Z.; Dorotkiewicz-Jach, A. Liposomes as Delivery Systems for Antibiotics. Int J Pharm 2010, 387 (1–2), 187–198. 10.1016/J.IJPHARM.2009.11.033.

31. Goodson, J. M. Antimicrobial Strategies for Treatment of Periodontal Diseases. Periodontol 2000 1994, 5 (1), 142–168. 10.1111/J.1600-0757.1994.TB00022.X.

32. Andersson, D. I.; Hughes, D. Microbiological Effects of Sublethal Levels of Antibiotics. Nature Reviews Microbiology 2014 12:7 2014, 12 (7), 465–478. 10.1038/nrmicro3270.

33. Kleinfelder, J. W.; Müller, R. F.; Lange, D. E. Antibiotic Susceptibility of Putative Periodontal Pathogens in Advanced Periodontitis Patients. J Clin Periodontol 1999, 26 (6), 347–351. 10.1034/J.1600-051X.1999.260603.X.

34. Kulik, E. M.; Thurnheer, T.; Karygianni, L.; Walter, C.; Sculean, A.; Eick, S. Antibiotic Susceptibility Patterns of *Aggregatibacter Actinomycetemcomitans* and *Porphyromonas Gingivalis* Strains from Different Decades. Antibiotics 2019, Vol. 8, Page 253 2019, 8 (4), 253. 10.3390/ANTIBIOTICS8040253.

35. Rams, T. E.; Degener, J. E.; van Winkelhoff, A. J. Antibiotic Resistance in Human Chronic Periodontitis Microbiota. J Periodontol 2014, 85 (1), 160–169. 10.1902/JOP.2013.130142.

36. Ardila, C. M.; Bedoya-García, J. A. Antimicrobial Resistance of *Aggregatibacter Actinomycetemcomitans*, *Porphyromonas Gingivalis* and *Tannerella Forsythia* in Periodontitis Patients. J Glob Antimicrob Resist 2020, 22, 215–218. 10.1016/J.JGAR.2020.02.024.

37. Walker, C. B. The Acquisition of Antibiotic Resistance in the Periodontal Microflora. Periodontol 2000 1996, 10 (1), 79–88. 10.1111/J.1600-0757.1996.TB00069.X.

38. Taichman, N. S.; Simpson, D. L.; Sakurada, S.; Cranfield, M.; DiRienzo, J.; Slots, J. Comparative Studies on the Biology of *Actinobacillus Actinomycetemcomitans* Leukotoxin in Primates. Oral Microbiol Immunol 1987, 2 (3), 97–104. 10.1111/J.1399-302X.1987.TB00270.X.

39. Lally, E. T.; Hill, R. B.; Kieba, I. R.; Korostoff, J. The Interaction between RTX Toxins and Target Cells. Trends Microbiol 1999, 7 (9), 356–361. 10.1016/S0966-842X(99)01530-9.

40. Johansson, A. *Aggregatibacter Actinomycetemcomitans* Leukotoxin: A Powerful Tool with Capacity to Cause Imbalance in the Host Inflammatory Response. Toxins 2011, Vol. 3, Pages 242-259 2011, 3 (3), 242–259. 10.3390/TOXINS3030242.

41. Åberg, C. H.; Kelk, P.; Johansson, A. *Aggregatibacter Actinomycetemcomitans*: Virulence of Its Leukotoxin and Association with Aggressive Periodontitis. Virulence 2015, 6 (3), 188–195. 10.4161/21505594.2014.982428.

42. Venketaraman, V.; Lin, A. K.; Le, A.; Kachlany, S. C.; Connell, N. D.; Kaplan, J. B. Both Leukotoxin and Poly-N-Acetylglucosamine Surface Polysaccharide Protect *Aggregatibacter Actinomycetemcomitans* Cells from Macrophage Killing. Microb Pathog 2008, 45 (3), 173–180. 10.1016/J.MICPATH.2008.05.007.

43. Haubek, D.; Johansson, A. Pathogenicity of the Highly Leukotoxic JP2 Clone of *Aggregatibacter Actinomycetemcomitans* and Its Geographic Dissemination and Role in Aggressive Periodontitis. J Oral Microbiol 2014, 6 (1). 10.3402/JOM.V6.23980.

44. Kachlany, S. C.; Fine, D. H.; Figurski, D. H. Secretion of RTX Leukotoxin by *Actinobacillus Actinomycetemcomitans*. Infect Immun 2000, 68 (11), 6094–6100. 10.1128/IAI.68.11.6094-6100.2000.

45. Linhartová, I.; Bumba, L.; Mašn, J.; Basler, M.; Osička, R.; Kamanová, J.; Procházková, K.; Adkins, I.; HejnováHolubová, J.; Sadílková, L.; Morová, J.; Šebo, P. RTX Proteins: A Highly Diverse Family Secreted by a Common Mechanism. FEMS Microbiol Rev 2010, 34 (6), 1076–1112. 10.1111/J.1574-6976.2010.00231.X.

46. Lally, E. T.; Kieba, I. R.; Sato, A.; Green, C. L.; Rosenbloom, J.; Korostoff, J.; Wang, J. F.; Shenker, B. J.; Ortlepp, S.; Robinson, M. K.; Billings, P. C. RTX Toxins Recognize a β2 Integrin on the Surface of Human Target Cells. Journal of Biological Chemistry 1997, 272 (48), 30463–30469. 10.1074/JBC.272.48.30463.

47. Brown, A. C.; Balashova, N. V.; Epand, R. M.; Epand, R. F.; Bragin, A.; Kachlany, S. C.; Walters, M. J.; Du, Y.; Boesze-Battaglia, K.; Lally, E. T. *Aggregatibacter Actinomycetemcomitans* Leukotoxin Utilizes a Cholesterol Recognition/Amino Acid Consensus Site for Membrane Association. Journal of Biological Chemistry 2013, 288 (32), 23607–23621. 10.1074/JBC.M113.486654.

48. Brown, A. C.; Boesze-Battaglia, K.; Du, Y.; Stefano, F. P.; Kieba, I. R.; Epand, R. F.; Kakalis, L.; Yeagle, P. L.; Epand, R. M.; Lally, E. T. *Aggregatibacter Actinomycetemcomitans* Leukotoxin Cytotoxicity Occurs through Bilayer Destabilization. Cell Microbiol 2012, 14 (6), 869–881. 10.1111/J.1462-5822.2012.01762.X.

49. Yeagle, P. L.; Sen, A. Hydration and the Lamellar to Hexagonal II Phase Transition of Phosphatidylethanolamine. Biochemistry 1986, 25 (23), 7518–7522. 10.1021/BI00371A039.

50. Koufos, E.; Chang, E. H.; Rasti, E. S.; Krueger, E.; Brown, A. C. Use of a Cholesterol Recognition Amino Acid Consensus Peptide to Inhibit Binding of a Bacterial Toxin to Cholesterol. Biochemistry 2016, 55 (34), 4787–4797. 10.1021/ACS.BIOCHEM.6B00430.

51. Krueger, E.; Hayes, S.; Chang, E. H.; Yutuc, S.; Brown, A. C. Receptor-Based Peptides for Inhibition of Leukotoxin Activity. ACS Infect Dis 2018, 4 (7), 1073–1081. 10.1021/acsinfecdis.7b00230.

52. Chang, E. H.; Huang, J.; Lin, Z.; Brown, A. C. Catechin-Mediated Restructuring of a Bacterial Toxin Inhibits Activity. Biochim Biophys Acta Gen Subj 2019, 1863 (1), 191– 198. 10.1016/j.bbagen.2018.10.011.

53. Chang, E. H.; Brown, A. C. Epigallocatechin Gallate Alters Leukotoxin Secretion and *Aggregatibacter Actinomycetemcomitans* Virulence. Journal of Pharmacy and Pharmacology 2021, 73 (4), 505–514. 10.1093/jpp/rgaa051.

54. van Gorkom, L. C. M.; Nie, S. Q.; Epand, R. M. Hydrophobic Lipid Additives Affect Membrane Stability and Phase Behavior of N- Monomethyldioleoylphosphatidylethanolamine. Biochemistry 1992, 31 (3), 671–677. 10.1021/BI00118A006.

55. Müller, H. P.; Holderrieth, S.; Burkhardt, U.; Höffler, U. In Vitro Antimicrobial Susceptibility of Oral Strains of *Actinobacillus Actinomycetemcomitans* to Seven Antibiotics. J Clin Periodontol 2002, 29 (8), 736–742. 10.1034/J.1600-051X.2002.290810.X.

56. Balashova, N. V.; Crosby, J. A.; Al Ghofaily, L.; Kachlany, S. C. Leukotoxin Confers Beta-Hemolytic Activity to *Actinobacillus Actinomycetemcomitans*. Infect Immun 2006, 74 (4), 2015–2021. 10.1128/iai.74.4.2015-2021.2006.

57. Teughels, W.; Dhondt, R.; Dekeyser, C.; Quirynen, M. Treatment of Aggressive Periodontitis. Periodontol 2000 2014, 65 (1), 107–133. 10.1111/PRD.12020.

58. Graziani, F.; Karapetsa, D.; Alonso, B.; Herrera, D. Nonsurgical and Surgical Treatment of Periodontitis: How Many Options for One Disease? Periodontol 2000 2017, 75 (1), 152–188. 10.1111/PRD.12201.

59. Greenstein, G. Clinical Significance of Bacterial Resistance to Tetracyclines in the Treatment of Periodontal Diseases. J Periodontol 1995, 66 (11), 925–932. 10.1902/JOP.1995.66.11.925.

60. Pinto-Alphandary, H.; Andremont, A.; Couvreur, P. Targeted Delivery of Antibiotics Using Liposomes and Nanoparticles: Research and Applications. Int J Antimicrob Agents 2000, 13 (3), 155–168. 10.1016/S0924-8579(99)00121-1.

61. Zambon, J. J.; Slots, J.; Genco, R. J. Serology of Oral *Actinobacillus Actinomycetemcomitans* and Serotype Distribution in Human Periodontal Disease. Infect Immun 1983, 41 (1), 19–27. 10.1128/IAI.41.1.19-27.1983.

62. Haubek, D.; Ennibi, O. K.; Poulsen, K.; Væth, M.; Poulsen, S.; Kilian, M. Risk of Aggressive Periodontitis in Adolescent Carriers of the JP2 Clone of *Aggregatibacter* (*Actinobacillus*) *Actinomycetemcomitans* in Morocco: A Prospective Longitudinal Cohort Study. The Lancet 2008, 371 (9608), 237–242. 10.1016/S0140-6736(08)60135-X.

63. Haraszthy, V. I.; Hariharan, G.; Tinoco, E. M. B.; Cortelli, J. R.; Lally, E. T.; Davis, E.; Zambon, J. J. Evidence for the Role of Highly Leukotoxic *Actinobacillus Actinomycetemcomitans* in the Pathogenesis of Localized Juvenile and Other Forms of Early-Onset Periodontitis. J Periodontol 2000, 71 (6), 912–922. 10.1902/JOP.2000.71.6.912.

64. Ferreira, M.; Ogren, M.; Dias, J. N. R.; Silva, M.; Gil, S.; Tavares, L.; Aires-Da-silva, F.; Gaspar, M. M.; Aguiar, S. I. Liposomes as Antibiotic Delivery Systems: A Promising Nanotechnological Strategy against Antimicrobial Resistance. Molecules 2021, 26 (7). 10.3390/molecules26072047.

65. Lee, Y.; Thompson, D. H. Stimuli-Responsive Liposomes for Drug Delivery. Wiley Interdiscip Rev Nanomed Nanobiotechnol 2017, 9 (5), e1450. 10.1002/WNAN.1450.

66. Zangabad, P. S.; Mirkiani, S.; Shahsavari, S.; Masoudi, B.; Masroor, M.; Hamed, H.; Jafari, Z.; Taghipour, Y. D.; Hashemi, H.; Karimi, M.; Hamblin, M. R. Stimulus-Responsive Liposomes as Smart Nanoplatforms for Drug Delivery Applications. Nanotechnol Rev 2018, 7 (1), 95–122. 10.1515/NTREV-2017-0154/ASSET/GRAPHIC/J_NTREV-2017-0154_FIG_005.JPG.

67. Meesters, C.; Brack, A.; Hellmann, N.; Decker, H. Structural Characterization of the α- Hemolysin Monomer from Staphylococcus Aureus. Proteins: Structure, Function, and Bioinformatics 2009, 75 (1), 118–126. 10.1002/PROT.22227.

68. Song, L.; Hobaugh, M. R.; Shustak, C.; Cheley, S.; Bayley, H.; Gouaux, J. E. Structure of Staphylococcal α-Hemolysin, a Heptameric Transmembrane Pore. Science (1979) 1996, 274 (5294), 1859–1866. 10.1126/SCIENCE.274.5294.1859.

69. Pornpattananangkul, D.; Zhang, L.; Olson, S.; Aryal, S.; Obonyo, M.; Vecchio, K.; Huang, C.-M.; Zhang, L. Bacterial Toxin-Triggered Drug Release from Gold Nanoparticle-Stabilized Liposomes for the Treatment of Bacterial Infection. J. Am. Chem. Soc 2011, 133, 4132–4139. 10.1021/ja111110e.

70. Yang, S.; Han, X.; Yang, Y.; Qiao, H.; Yu, Z.; Liu, Y.; Wang, J.; Tang, T. Bacteria-Targeting Nanoparticles with Microenvironment-Responsive Antibiotic Release to Eliminate Intracellular Staphylococcus Aureus and Associated Infection. ACS Appl Mater Interfaces 2018, 10 (17), 14299–14311. 10.1021/ACSAMI.7B15678/ASSET/IMAGES/LARGE/AM-2017-156786_0009.JPEG.

71. Ludwig, A.; Goebel, W. Structure and Mode of Action of RTX Toxins. The Comprehensive Sourcebook of Bacterial Protein Toxins 2006, 547–569. 10.1016/B978-012088445-2/50034-2.

72. Linhartová, I.; Bumba, L.; Mašn, J.; Basler, M.; Osička, R.; Kamanová, J.; Procházková, K.; Adkins, I.; HejnováHolubová, J.; Sadílková, L.; Morová, J.; Šebo, P. RTX Proteins: A Highly Diverse Family Secreted by a Common Mechanism. FEMS Microbiol Rev 2010, 34 (6), 1076–1112. 10.1111/J.1574-6976.2010.00231.X.

73. Bakás, L.; Chanturiya, A.; Herlax, V.; Zimmerberg, J. Paradoxical Lipid Dependence of Pores Formed by the *Escherichia Coli* α-Hemolysin in Planar Phospholipid Bilayer Membranes. Biophys J 2006, 91 (10), 3748–3755. 10.1529/BIOPHYSJ.106.090019.

74. Zitzer, A.; Bittman, R.; Verbicky, C. A.; Erukulla, R. K.; Bhakdi, S.; Weis, S.; Valeva, A.; Palmer, M. Coupling of Cholesterol and Cone-Shaped Lipids in Bilayers Augments Membrane Permeabilization by the Cholesterol-Specific Toxins Streptolysin O and *Vibrio Cholerae* Cytolysin. Journal of Biological Chemistry 2001, 276 (18), 14628–14633. 10.1074/jbc.M100241200.

75. Martín, C.; Requero, M. A.; Masin, J.; Konopasek, I.; Goñi, F. M.; Sebo, P.; Ostolaza, H. Membrane Restructuring by *Bordetella Pertussis* Adenylate Cyclase Toxin, a Member of the RTX Toxin Family. J Bacteriol 2004, 186 (12), 3760–3765. 10.1128/JB.186.12.3760-3765.2004.

76. Ostolaza, H.; Bartolomé, B.; de Zárate, I. O.; de la Cruz, F.; Goñi, F. M. Release of Lipid Vesicle Contents by the Bacterial Protein Toxin α-Haemolysin. Biochimica et Biophysica Acta (BBA) - Biomembranes 1993, 1147 (1), 81–88. 10.1016/0005-2736(93)90318-T.

77. Amuategi, J.; Alonso, R.; Ostolaza, H. Four Cholesterol-Recognition Motifs in the Pore-Forming and Translocation Domains of Adenylate Cyclase Toxin Are Essential for Invasion of Eukaryotic Cells and Lysis of Erythrocytes. International Journal of Molecular Sciences 2022, Vol. 23, Page 8703 2022, 23 (15), 8703. 10.3390/IJMS23158703.

78. Osickova, A.; Balashova, N.; Masin, J.; Sulc, M.; Roderova, J.; Wald, T.; Brown, A. C.; Koufos, E.; Chang, E. H.; Giannakakis, A.; Lally, E. T.; Osicka, R. Cytotoxic Activity of *Kingella Kingae* RtxA Toxin Depends on Post-Translational Acylation of Lysine Residues and Cholesterol Binding. Emerg Microbes Infect 2018, 7 (1). 10.1038/S41426-018-0179-X.

79. González Bullón, D.; Uribe, K. B.; Amuategi, J.; Martín, C.; Ostolaza, H. Cholesterol Stimulates the Lytic Activity of Adenylate Cyclase Toxin on Lipid Membranes by Promoting Toxin Oligomerization and Formation of Pores with a Greater Effective Size. FEBS Journal 2021, 288 (23), 6795–6814. 10.1111/FEBS.16107.

80. Henry, B. D.; Neill, D. R.; Becker, K. A.; Gore, S.; Bricio-Moreno, L.; Ziobro, R.; Edwards, M. J.; Mühlemann, K.; Steinmann, J.; Kleuser, B.; Japtok, L.; Luginbühl, M.; Wolfmeier, H.; Scherag, A.; Gulbins, E.; Kadioglu, A.; Draeger, A.; Babiychuk, E. B. Engineered Liposomes Sequester Bacterial Exotoxins and Protect from Severe Invasive Infections in Mice. Nat Biotechnol 2015, 33 (1), 81–88. 10.1038/nbt.3037.

81. Fong, K. P.; Pacheco, C. M. F.; Otis, L. L.; Baranwal, S.; Kieba, I. R.; Harrison, G.; Hersh, E. V.; Boesze-Battaglia, K.; Lally, E. T. *Actinobacillus Actinomycetemcomitans* Leukotoxin Requires Lipid Microdomains for Target Cell Cytotoxicity. Cell Microbiol 2006, 8 (11), 1753–1767. 10.1111/J.1462-5822.2006.00746.X.

82. Hu, C.-M. J.; Fang, R. H.; Copp, J.; Luk, B. T.; Zhang, L. A Biomimetic Nanosponge That Absorbs Pore-Forming Toxins. NATURE NANOTECHNOLOGY | 2013, 8. 10.1038/NNANO.2013.54.

83. Jiang, L.; Zhu, Y.; Luan, P.; Xu, J.; Ru, G.; Fu, J. G.; Sang, N.; Xiong, Y.; He, Y.; Lin, G. Q.; Wang, J.; Zhang, J.; Li, R. Bacteria -Anchoring Hybrid Liposome Capable of Absorbing Multiple Toxins for Antivirulence Therapy of *Escherichia Coli* Infection. ACS Nano 2021, 15 (3), 4173–4185. 10.1021/ACSNANO.0C04800/ASSET/IMAGES/LARGE/NN0C04800_0006.JPEG.

84. Henry, B. D.; Neill, D. R.; Becker, K. A.; Gore, S.; Bricio-Moreno, L.; Ziobro, R.; Edwards, M. J.; Mühlemann, K.; Steinmann, J.; Kleuser, B.; Japtok, L.; Luginbühl, M.; Wolfmeier, H.; Scherag, A.; Gulbins, E.; Kadioglu, A.; Draeger, A.; Babiychuk, E. B. Engineered Liposomes Sequester Bacterial Exotoxins and Protect from Severe Invasive Infections in Mice. Nature Biotechnology 2014 33:1 2014, 33 (1), 81–88. 10.1038/nbt.3037.

85. Breijyeh, Z.; Jubeh, B.; Karaman, R. Resistance of Gram-Negative Bacteria to Current Antibacterial Agents and Approaches to Resolve It. Molecules 2020, Vol. 25, Page 1340 2020, 25 (6), 1340. 10.3390/MOLECULES25061340.

86. Tsuzukibashi, O.; Takada, K.; Saito, M.; Kimura, C.; Yoshikawa, T.; Makimura, M.; Hirasawa, M. A Novel Selective Medium for Isolation of *Aggregatibacter* (*Actinobacillus*) *Actinomycetemcomitans*. J Periodontal Res 2008, 43 (5), 544–548. 10.1111/j.1600-0765.2007.01074.x.

87. Balashova, N. v.; Shah, C.; Patel, J. K.; Megalla, S.; Kachlany, S. C. *Aggregatibacter Actinomycetemcomitans* LtxC Is Required for Leukotoxin Activity and Initial Interaction between Toxin and Host Cells. Gene 2009, 443 (1–2), 42–47. 10.1016/j.gene.2009.05.002.

88. Lally, E. T.; Golub, E. E.; Kieba, I. R. Identification and Immunological Characterization of the Domain of *Actinobacillus Actinomycetemcomitans* Leukotoxin That Determines Its Specificity for Human Target Cells. Journal of Biological Chemistry 1994, 269 (49), 31289–31295. 10.1016/S0021-9258(18)47421-2.

89. Schneider, C. A.; Rasband, W. S.; Eliceiri, K. W. NIH Image to ImageJ: 25 Years of Image Analysis. Nat Methods 2012, 9 (7), 671. 10.1038/NMETH.2089.

90. Abramoff, M. D.; Magalhães, P. J.; Ram, S. J. Image Processing with ImageJ. Biophotonics international 2004, 11 (7), 36–42.

