## Supporting Information for "Toxin-Triggered Liposomes for the Controlled Release of Antibiotics to Treat Infections Associated with Gram-Negative Bacteria"

**Development of a Toxin-Triggered Liposomal Antibiotic Delivery Vehicle for the Treatment of *Aggregatibacter actinomycetemcomitans***

**Ziang Li^1^, Rani Baidoun^1,2^, Angela C. Brown^1*^**

^1^Department of Chemical and Biomolecular Engineering, Lehigh University, Bethlehem, PA

^2^Current Affiliation: Department of Chemical and Biomolecular Engineering, University of Pennsylvania, Philadelphia, PA

**Table S1. Statistical analysis of the LtxA-mediated leakage data shown in Figure 2B**

| **ANOVA Results** | | | | |
| --- | --- | --- | --- | --- |
| **Analyzed groups** | | | **p-value** | **Significance** |
| 100% N-methyl-DOPE | vs | 80% N-methyl-DOPE/20% chol | 1.07 x 10^-1^ | NS |
| 100% N-methyl-DOPE | vs | 60% N-methyl-DOPE/40% chol | 7.53 x 10^-5^ | **** |
| 80% N-methyl-DOPE/20% chol | vs | 60% N-methyl-DOPE/40% chol | 2.28 x 10^-3^ | *** |
| 60% DOPC/40% chol | vs | 60% N-methyl-DOPE/40% chol | 6.85 x 10^-7^ | **** |

NS, not significant; ***, p ≤ 0.005; ****, p ≤ 0.001

**Table S2. Mean hydrodynamic radius and PDI of empty and moxifloxacin-loaded liposomes stored at 4 °C or 22 °C over 7 days**

|  | **Empty Liposomes** | | | | **Moxifloxacin-Loaded Liposomes** | | | |
| --- | --- | --- | --- | --- | --- | --- | --- | --- |
|  | **Storage at 4 °C** | | **Storage at 22 °C** | | **Storage at 4 °C** | | **Storage at 22 °C** | |
| **Day** | **Diameter**  **(nm)** | **PDI** | **Diameter**  **(nm)** | **PDI** | **Diameter**  **(nm)** | **PDI** | **Diameter**  **(nm)** | **PDI** |
| 0 | 196 | 0.222 | 194 | 0.117 | 152 | 0.103 | 200 | 0.102 |
| 1 | 202 | 0.237 | 170 | 0.150 | 157 | 0.127 | 186 | 0.087 |
| 2 | 215 | 0.255 | 155 | 0.125 | 169 | 0.136 | 219 | 0.072 |
| 3 | 210 | 0.243 | 226 | 0.132 | 187 | 0.114 | 206 | 0.109 |
| 4 | 220 | 0.180 | 175 | 0.153 | 218 | 0.115 | 211 | 0.104 |
| 5 | 206 | 0.290 | 209 | 0.201 | 203 | 0.180 | 198 | 0.155 |
| 6 | 206 | 0.290 | 219 | 0.177 | 191 | 0.154 | 214 | 0.136 |
| 7 | 230 | 0.200 | 238 | 0.142 | 198 | 0.205 | 220 | 0.141 |

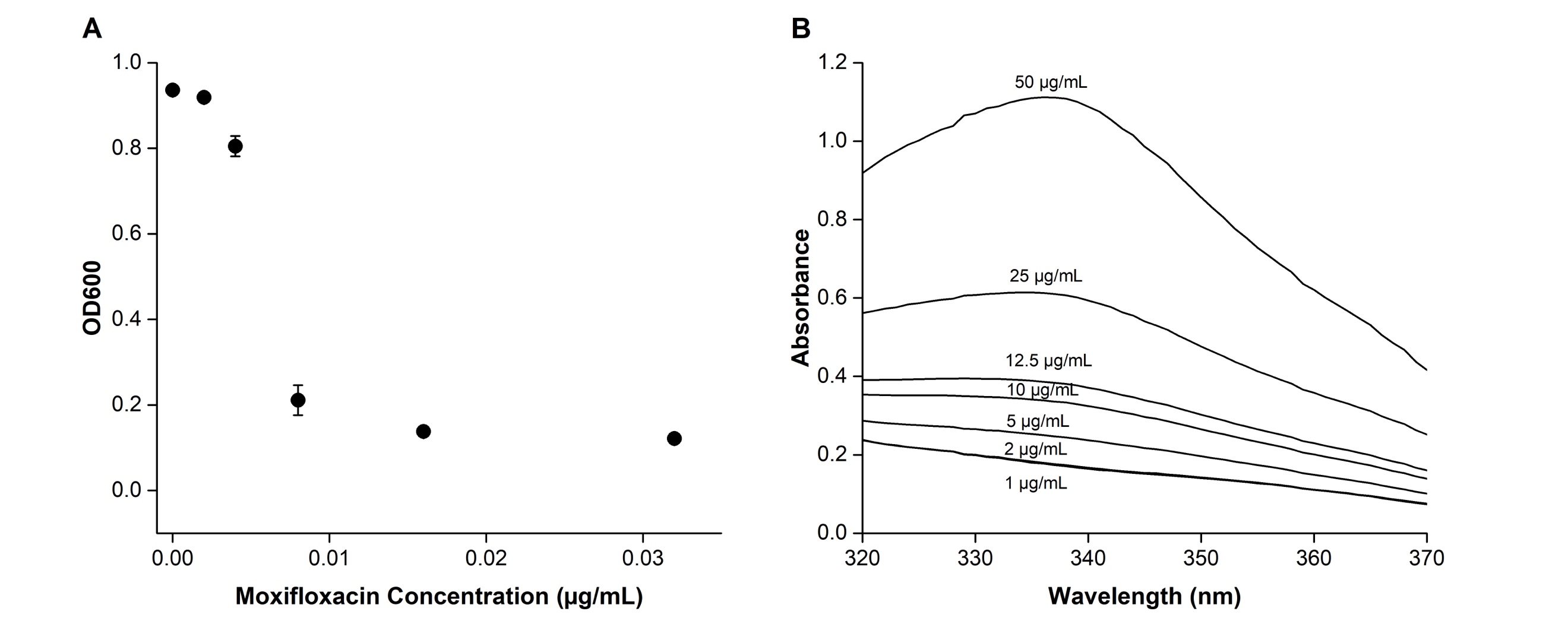

**Figure S1. Moxifloxacin Efficacy and Encapsulation.** (A) Moxifloxacin Efficacy. *A. actinomycetemcomitans* JP2 cells were grown for 24 hours in the presence of moxifloxacin at different concentrations. OD_600_ readings were collected in an UV-vis spectrophotometer. (B) UV-vis absorption spectra of moxifloxacin. The absorbance of moxifloxacin at different concentrations was scanned from 320 nm to 370 nm, and the absorption maximum was observed to be at a wavelength of approximately 335 nm.

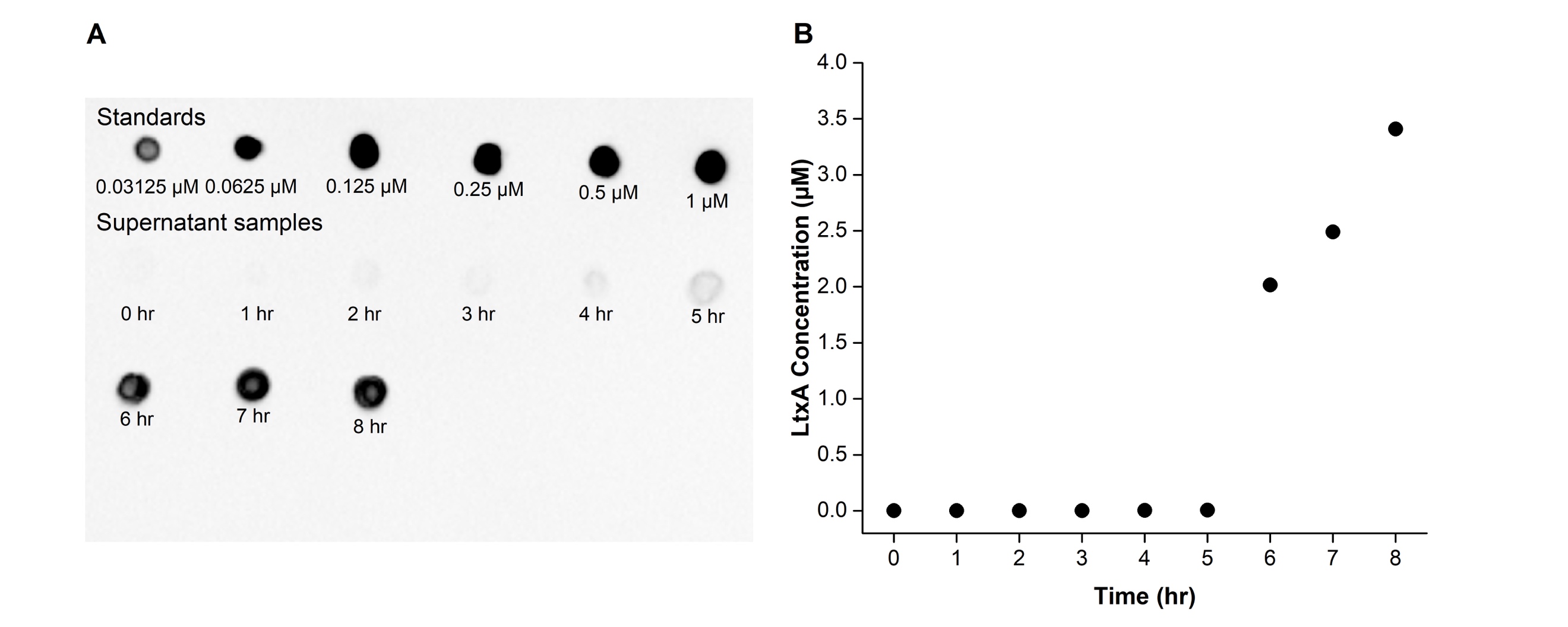

**Figure S2. Timing of LtxA production.** (A) Immunoblot of LtxA in the *A. actinomycetemcomitans* JP2 cell culture supernatant at varying culture times (bottom rows) compared to LtxA standards at known concentrations (top row). A calibration curve was created using the intensity of each dot measured in ImageJ and the LtxA concentrations of the standards, which was then used to determine the LtxA concentration in the *A. actinomycetemcomitans* supernatants collected at different time points. (B) LtxA production kinetics over 8 hours. The concentration of LtxA in the *A. actinomycetemcomitans* supernatant was plotted as a function of time, indicating that LtxA production was initiated after six hours of growth.

**Table S3. Statistical analysis of the moxifloxacin-loaded liposome efficacy data shown in Figure 5**

| **ANOVA Results** | | | | |
| --- | --- | --- | --- | --- |
| **Analyzed groups** | | | **p-value** | **Significance** |
| JP2, No treatment | vs | JP2, Mox-loaded liposomes | 0.02 | * |
| AA1704, No treatment | vs | AA1704, Mox-loaded liposomes | 0.20 | NS |
| JP2, Mox-loaded liposomes | vs | AA1704, Mox-loaded liposomes | 0.01 | ** |

NS, not significant; *, p ≤ 0.05; **, p ≤ 0.01
